# Inverse hourglass pattern of conservation in rodent molar development

**DOI:** 10.1101/2025.01.23.634446

**Authors:** Jérémy Ganofsky, Mathilde Estevez-Villar, Marion Mouginot, Sébastien Moretti, Marion Nyamari, Marc Robinson-Rechavi, Sophie Pantalacci, Marie Sémon

## Abstract

Although it is well established that certain stages of development are more conserved than others, the reasons for this phenomenon remain largely unknown. We study molecular conservation in the development of an organ, the molar, by comparing the temporal profiles of expression in mice and hamsters. We find that molar development is characterized by a rarely observed pattern of conservation of expression level and coding sequences forming an inverse hourglass, with more conservation at the beginning and end of morphogenesis than at intermediate stages. As the development of the rodent molar is well described, we were able to link this pattern to the properties of the expressed genes and the activity of different developmental processes. Early and late stages mobilize different sets of pleiotropic genes, cell division for bud growth and secretion for tooth mineralization. The particularities of dental morphogenesis and homeostasis, with the degradation of certain tissues at the end of development and the hosting of immune cells, as well as heterochronies linked to adaptations, also probably contribute to the pattern. Our study of the different actors explains the inverted hourglass of molars by a combination of processes intrinsic to the teeth, and by negative and positive selection which is mostly extrinsic to the teeth. This is likely translatable to explain molecular conservation patterns in many other biological systems.

## INTRODUCTION

Comparisons of embryos from different species, first in terms of morphology and more recently in terms of gene expression, show that certain stages are more conserved than others. For example, in many clades, embryonic development is particularly conserved at the phylotypic stage, when highly conserved key developmental genes orchestrate body plan organization [1]. Transcriptomics, which allows genome expression to be sampled and compared at different developmental stages between species, has become an effective quantitative approach to measure embryonic molecular conservation [2–7].

Molecular conservation has been measured from transcriptomics in different ways, either directly as the conservation of gene expression levels across stages, or indirectly as properties of the genes expressed, such as protein sequence conservation or gene phylogenetic age. In the later case, this can be done by establishing groups of stage-specific genes, or by weighting or correlating different properties and gene expression levels. In most cases, these measurements have revealed patterns of differential conservation across stages e.g. [8–10].

Most studies have focused on molecular conservation in the whole developing embryo, and revealed an hourglass pattern at the phylotypic stage, followed by increasing molecular divergence in later stages [2,3,6–8,11,12].

Beyond describing patterns, much work has been devoted to deciphering the evolutionary processes that create them, and their relation to environmental adaptations or internal causes, such as development or genetic properties. It was proposed that very early and late stages are more subject to positive selection linked to ecological adaptations [13]. It has also been argued that the parts of an embryo or organ are initially interdependent, while this diminishes during development [1]. Therefore, mutations in genes expressed early in development tend to have major effects and should often be subject to purifying selection. Such hallmarks of purifying and positive selection were detected on coding and regulatory sequences (Garfield and Wray 2009; Liu et al. 2021). The importance of internal genetic constraints such as pleiotropy has also long been emphasized [1,4,8]. In functional genomics studies, pleiotropy is operationally defined by expression in a diversity of organs, developmental stages, or cell types. Thus defined pleiotropic genes are highly mobilized during early development, which is thought to reduce the number of mutations available for positive selection. The effects of functional constraints and adaptive changes are nonexclusive [8,14,15].

Molecular divergence in the development of individual organs has been much less studied than in the whole embryo. One study, a large-scale analysis of gene expression in seven mammalian organs, confirmed that conservation declines at later stages of organogenesis [10]. Genes active in early organ development had a broader spatial and temporal expression than genes employed later. Such a decrease in pleiotropy can explain both a decrease in functional constraints and an increase in adaptation. Indeed, the extent of purifying selection gradually decreased during development, whereas the amount of positive selection and expression of new genes increased. Other studies focused on a single organ, such as terminal inflorescence in grasses [16], or limbs in vertebrates [17,18]. A comparison of the forelimb buds of the mouse (*Mus musculus*) with the pectoral fin buds of the brown-banded bamboo shark (*Chiloscyllium punctatum*), revealed a stronger conservation of gene expression at mid-limb development and many heterochronic changes [18]. They also found that at intermediate stages of development, limb buds expressed pleiotropic genes that have conserved stage– and tissue-specific enhancers. They hypothesized that this complex regulation was vulnerable to genetic mutations, limiting the evolvability of this particular period of morphogenesis. Overall, both studies highlighted that pleiotropy is a major contributor to the variation in molecular divergence observed between species during organ development.

To go further and interpret the patterns of molecular conservation in relation to the evolution of adult phenotypes and that of morphogenesis processes, we need more closely related species, and organs with well known evolution and development. Here we chose the upper and lower molars, in two rodent species, the golden hamster (*Mesocricetus auratus*) and the mouse (*M. musculus*), that diverged ≈26 million years ago [19]. Molar tooth development has been well described [20–23]. In the mouse lineage, the upper molar underwent drastic adaptive changes that have been linked to the evolution of the morphogenetic processes of both teeth [24]. Here we have extended a previously published transcriptome time series, and established patterns of conservation of expression as well as adaptive and purifying selection in the coding sequence throughout molar development.

## RESULTS

### Transcriptomic profiling reveals gene sets whose expression peaks at specific stages of molar morphogenesis

To compare the temporal dynamics of gene expression between mouse and hamster molar development, we gathered 103 bulk RNA-seq samples from a time series of growing upper and lower molar tooth germs (36 samples were collected specifically for this study to complete our previously published data, Table S1). We selected tooth germs from Embryonic day (E)12.5 to PostNatal day (PN)2 mice because this is the period during which the major events of morphogenesis occur, from bud, to cap, to bell stage until differentiation and enamel/dentin secretion. For the hamster, we selected the stages of dental development over a period as wide as in the mouse (from E11.0 to PN2). The global relationship among all samples was explored through a principal component analysis (PCA) (Fig 1A and S1). The first principal component (PC1), explaining most variation in gene expression, separated the samples by species, confirming the coevolution of upper and lower molar [24]. PC2 separated the samples by developmental age, while batch effect explains only 9% of the variation (Fig S1B).

**Fig 1.**
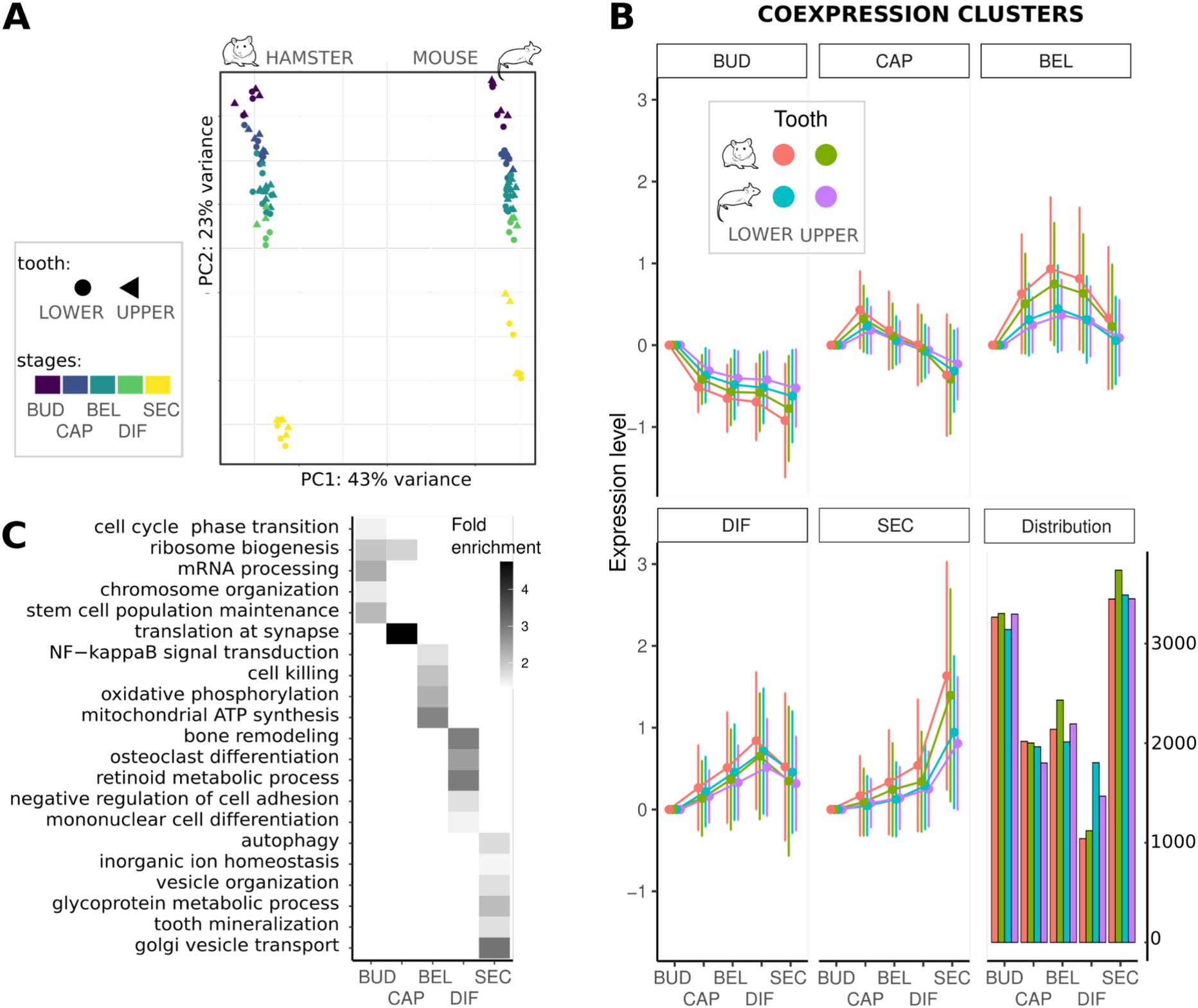
Global expression profiles and clusters of co-expressed genes with an expression peak at a specific stage of dental development. (*A) PCA based on 11,342 1:1 orthologs across mouse and hamster. Samples are colored per morphological stage, shapes correspond to upper and lower molar samples. (B) Coexpression clusters defined by soft clustering of expression values of each species and tooth type. Each cluster corresponds to a set of genes with an expression peak at a specific stage of dental development. The average profiles of each cluster are represented, with expression levels centered at the value of the first stage and error bars showing standard deviations. Clusters are ordered according to their temporal profiles. Distribution of cluster sizes is given. Genes not associated with a cluster are not represented. (C) Summary of the GO enrichment analysis for the mouse lower molar coexpression clusters; enriched GO terms were categorized as shown in Fig S2, a maximum of 5 representative terms are shown with their fold enrichments*.

To obtain the patterns of evolutionary conservation we first needed to group genes expressed at similar developmental stages. We attributed samples to five morphological stages: bud, cap, bell stage, differentiation and enamel secretion (Fig 1A, Table S1). Few genes are specifically expressed in a single stage (Table S2). Simple presence/absence calls are therefore not applicable to our data to study evolutionary conservation. We decided instead to classify genes according to quantitative variation in expression during development.

Applying a flexible clustering approach with semi-supervised starting conditions [25,26] to each species and tooth type, we clustered together genes whose expression peaked at the same stage. We call these “clusters of coexpression” further on.

We associated about three-quarters of the genes with a cluster, confirming the labile nature of gene expression during development, even in a relatively short period of morphogenesis and in a single organ. The distribution of cluster sizes is similar for all teeth, with more genes associated with early and late stages than with intermediate stages (Fig 1B).

To associate these clusters with biological functions, we performed a gene ontology (GO) enrichment analysis on each cluster (Fig 1C, S2, S3). In the mouse lower molar, we observed an initial phase associated with cell cycle, protein synthesis, and metabolism (bud), followed by an intermediate phase with similar (yet more modest) enrichments plus indicators of nerve colonization (cap), then by a phase associated with cell-cell signaling and mitochondrial respiration (bell), a phase associated with bone formation and blood cells (differentiation), and finally a phase associated with autophagy, secretion, and mineralisation (secretion). This temporal sequence is in agreement with the succeeding processes that contribute to molar formation [27]. Similar enrichment terms of GO terms were observed for the other molars.

### Shifts in expression peaks are less frequent for early or late peaking genes

Having clustered together genes whose expression peaked at the same stage in the time series of each tooth, we systematically compared the gene content of these clusters to determine whether the same genes peak at the same stage in each tooth. We measured their overlap in terms of enrichment (fold change) relative to random expectation (see Methods). The largest overlaps were observed between upper and lower molars of the same species (Fig S4), where clusters corresponding to the same stage have 60-70% of genes in common, which is 3 times more than would be obtained by chance. This is consistent with previous results on lower and upper molar transcriptome coevolution [24].

Although overlap was also observed between species, there were marked differences in gene content (Fig 2), again consistent with the rapid change in developmental expression observed in a previous study [24]. A third of the genes associated with a cluster whose expression peaks at a given stage in the upper molar of the mouse was associated with the cluster peaking at the same stage in the upper molar of the hamster, which is 50% more than expected (Fig 2C) but still a minority of the genes. The largest overlaps are observed for clusters with expression peaks at early and late stages. The difference is statistically significant: its magnitude far exceeds the bootstrap intervals (segments on the solid lines) and it exceeds the temporal variation observed on the shuffled data (dashed lines, Fig 2C,D). While higher similarity at the earliest stage was expected from previous studies of organ evolution, the high similarity of transcription profiles at late stages was more surprising. Thus we decided to explore this pattern in more detail at the level of sequence evolution.

**Fig 2.**
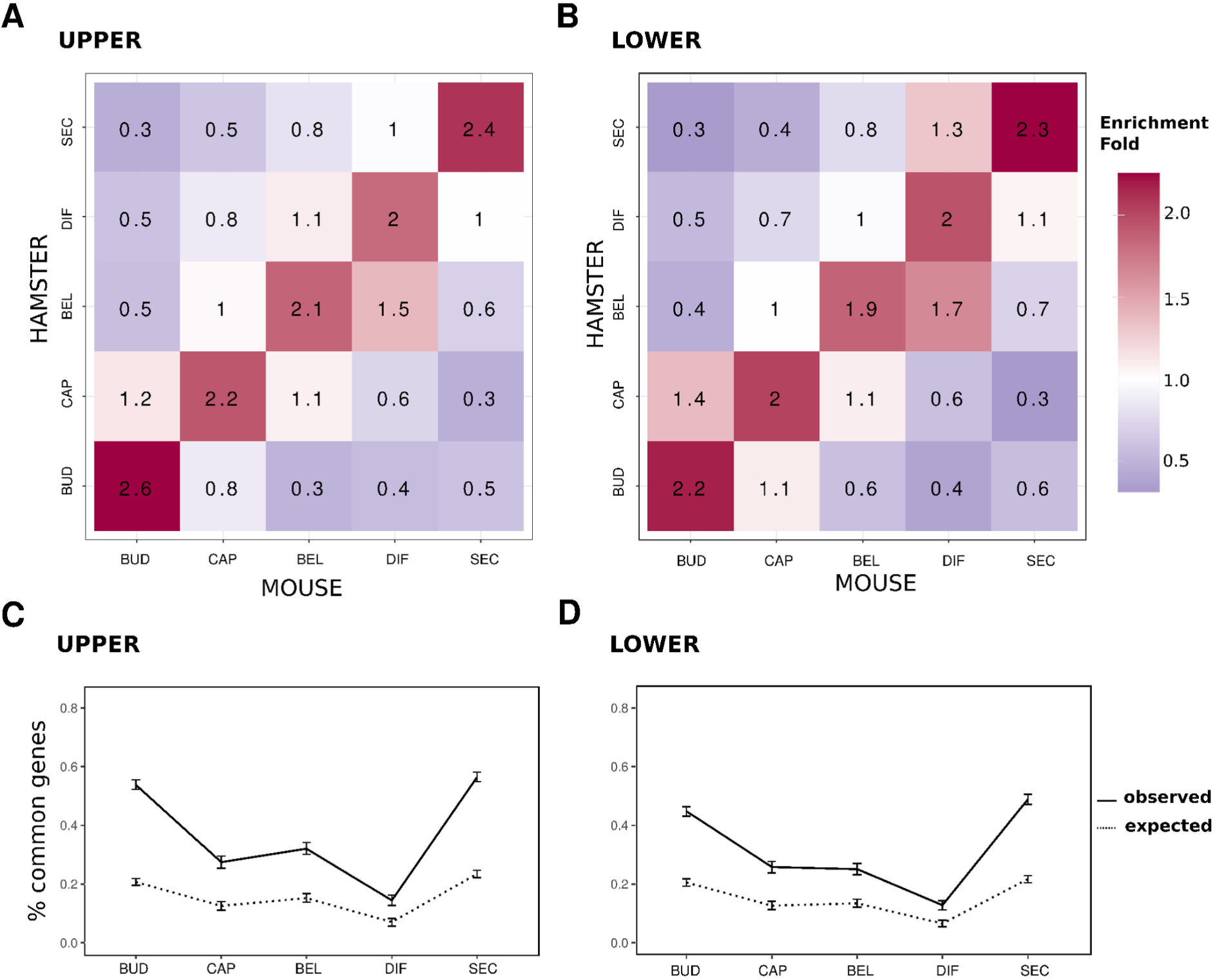
Overlap of expression gene clusters between species. *Each cluster corresponds to a set of genes with an expression peak at a specific stage of development in a given tooth and species. Number of genes shared between hamster and mouse coexpression clusters, expressed as a fold enrichment relative to equally sized random clusters, for the upper (A) and the lower (B) molars. Clusters are ordered according to their temporal profiles. Proportion of genes shared between hamster and mouse coexpression clusters peaking at the same stage in both species, in the upper (C) and lower (D) molars. Intervals show the 95 percentiles obtained by 1000 bootstrap resamplings. Expected values obtained by randomly shuffling genes between clusters (dashed lines)*.

### Constraints on coding sequences are higher for both genes expressed at early and at late stages

To assess potential differences in functional constraints during development, we compared the selective pressure operating on gene coding sequences between coexpressed gene clusters. We quantified the mode and strength of selection by using dN/dS ratios.

We obtained substitution rates per gene, and averaged them per coexpressed gene cluster for each tooth, to compare average values of dN/dS ratios between clusters associated with different stages. The average values of dN/dS ratios are well below one, indicating that purifying selection dominates coding sequence evolution in genes expressed in molars, as expected. What is more interesting is that ratios are significantly lower at early and late stages than at intermediate stages. This is true for all tooth types, with a maximum ratio (i.e. weakest constraint) for the differentiation stage in the mouse, and for the bell stage in the hamster. This creates an inverse hourglass pattern of purifying selection levels operating on coding sequences (Fig 3).

**Fig 3.**
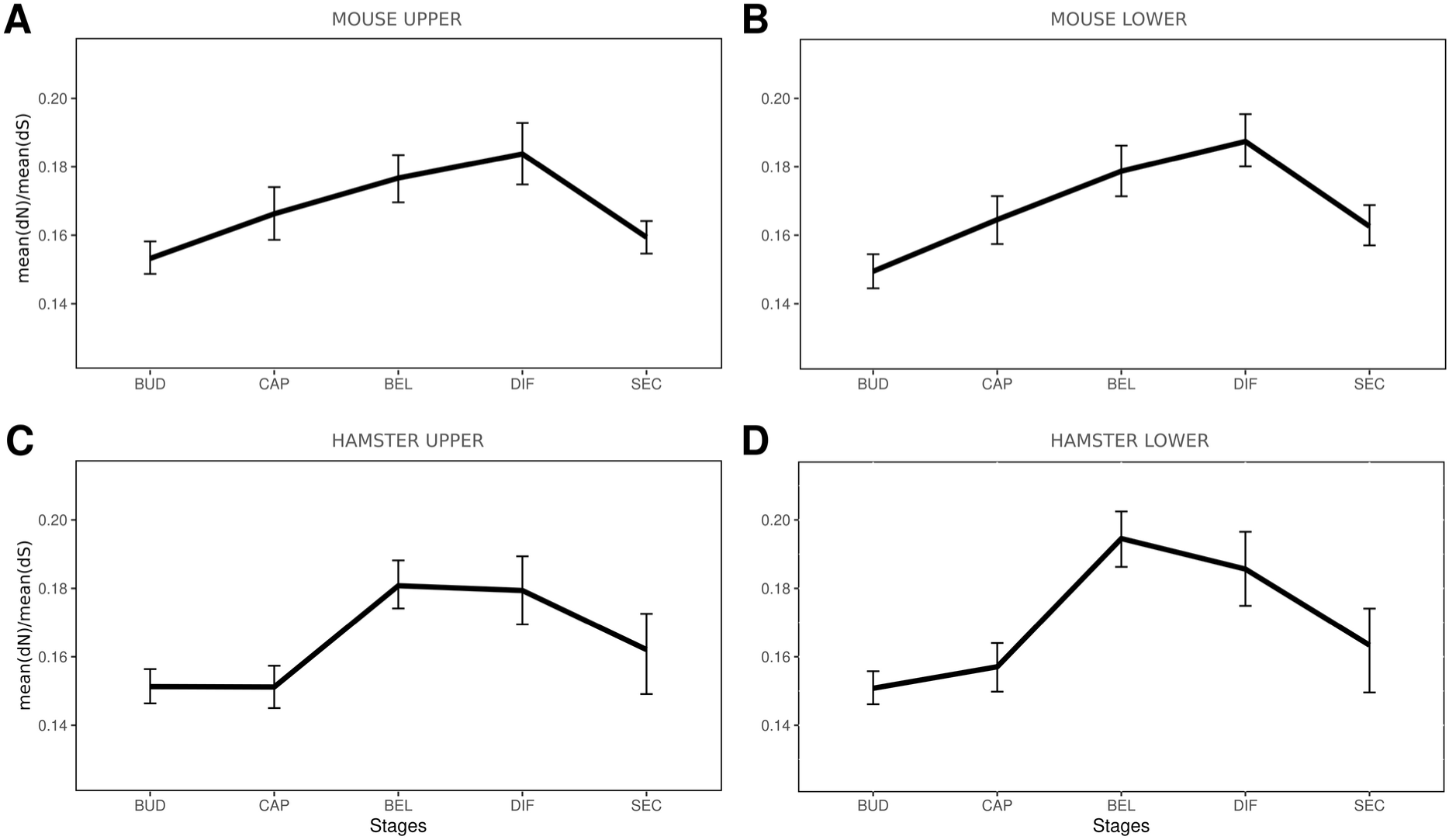
Selective pressure acting on gene coding sequences for each coexpressed gene cluster. *Average pairwise dN/dS between mouse and hamster were computed as mean(dN)/mean(dS). Observed values are represented by a black line, intervals show the 95 percentiles obtained by 1000 bootstrap resamplings*.

We verified this pattern both for genes with conserved and with non-conserved expression peaks between pairs of molars (Fig S5). In addition, genes with a conserved expression peak have significantly lower dN/dS values, indicating that conserved expression profiles are correlated with stronger purifying selection on their coding sequence. One possibility is that dN/dS profiles reflect different levels of constraints on tooth development. Another possibility is that this reflects the temporal usage of different sets of genes, whose dN/dS profiles are dictated by their role in many parts of the body, i.e. by their pleiotropy (see below).

### Adaptation in coding sequences tend to increase in mid molar development

Next, we evaluated the relationship between adaptation and development stages by examining the genes identified as carrying signatures of positive selection. We computed the proportion of genes with significant evidence of positive selection on several branches of the rodent tree (q-value < 0.2, Table S3). We found 18 genes on the *Muridae* branch (out of 11460 tested genes), 60 on the *Murinae* branch (12919 genes), 49 on the *Mus* branch (13257 genes), and 36 on the *Cricetinae* branch (10936 genes). Except for one gene, Csf1r, these genes have no clear role in tooth development [28], but many are involved in immunity (Fig S6A). This suggests that they were selected for other purposes than for tooth function, even though we cannot exclude that they may later serve immunity in the functional adult tooth. We grouped the genes by their associated stage and looked for pathway enrichments: bel/differentiation genes displayed significantly more interactions than expected with an enrichment for inflammatory response (PPI enrichment p-value, 0.019, N=47) while the genes associated with other stages did not (for bud/cap, N=42 genes and p-value=0.163, for secretion, N=30 and p-value=0.134, Fig S6B).

To compare the magnitude of positive selection across different stages, we used the likelihood ratio (Δln*L)* of the models with and without positive selection, following [15,29]. A branch in a gene tree with a higher Δln*L* value indicates higher evidence for positive selection for this gene over this branch. We computed Δln*L* for each gene and then took the average per coexpressed gene cluster (see Methods). The amount of positive selection increases from the early stages and is maximum at the bel/differentiation stages. This trend is particularly marked on the *Mus* branch but visible in all branches (Fig 4). Again, the lower evidence for positive selection at the earliest and last stages of development is consistent with the stronger evolutionary conservation seen in peak expression and in coding sequence purifying selection.

**Fig 4.**
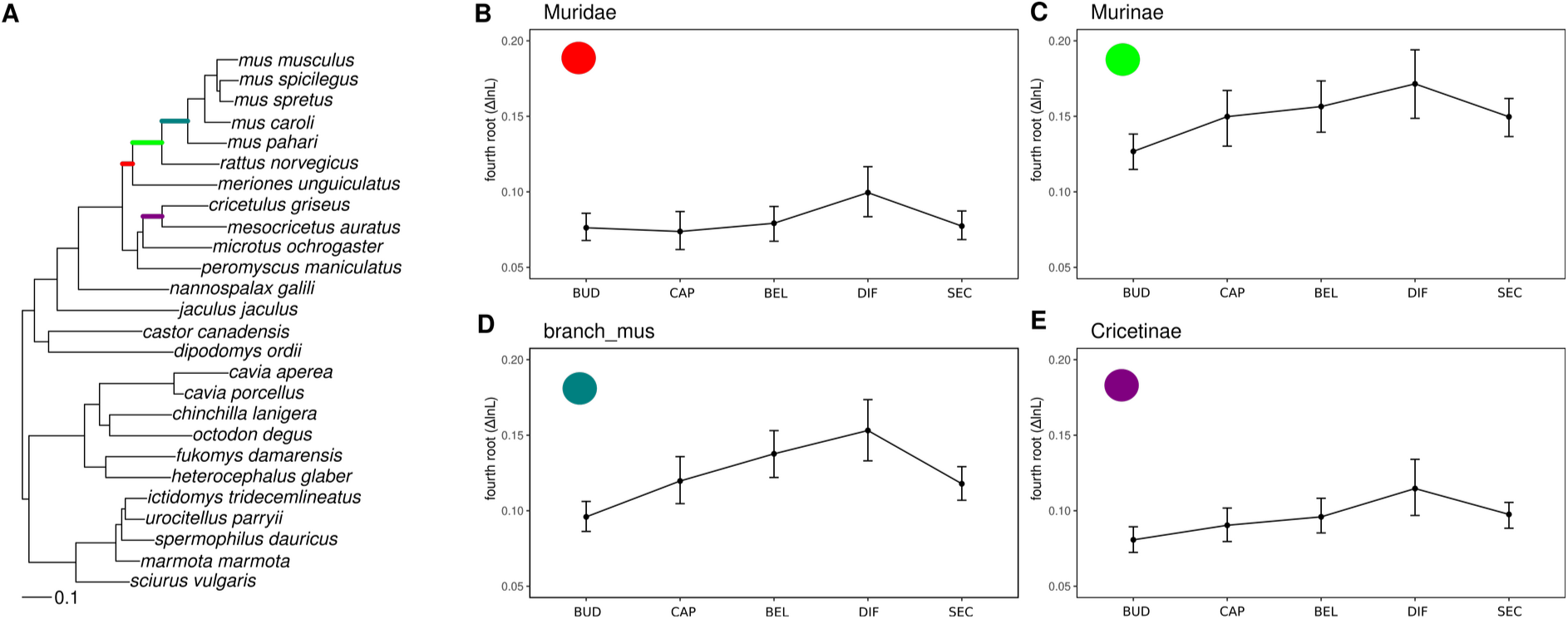
Positive selection for genes with expression peaks at different stages of development, in four branches of the rodent tree. *Each plot shows the fourth root of the likelihood ratio (ΔlnL) of the models with and without positive selection, taken from Selectome database. Average for coexpressed gene clusters associated with each stage of development are represented. Confidence intervals obtained from 1000 bootstraps. Branches used to compute positive selection are indicated on the rodent tree in (A), and correspond respectively to (B): Muridae, (C): Murinae, (D): Mus, (E): Cricetinae. Gene clusters correspond to mouse upper molar (B-D) or to hamster upper molar (E). Species tree in panel A pruned from Ensembl Compara*.

### Variation in expression levels and pleiotropy explain the developmental pattern of sequence conservation

The influence of different parameters such as levels and breath of expression on the rates of protein coding sequence evolution has been extensively studied in many organisms [30–34]. In mouse, genes with high expression levels tend to evolve under strong purifying selection, although the effect size is small when confounding factors are taken into account [30]. We observed this negative correlation in our data, indicating that expression in molar development correlates with the intensity of purifying selection on the coding sequence of genes (Fig S7).

We next tested whether the relationship between dN/dS and expression level impacted the patterns we observed in Fig 3. We split genes in quantiles according to their maximum expression level and computed mean dN/dS separately for each expression category. We confirmed that dN/dS was higher for genes in low expression quantiles than for genes in high expression quantiles (Fig S8). Within expression quantiles, there was a slight variation in dN/dS over the different developmental stages, but it was not significant. Hence the “inverse hourglass” pattern of conservation is seen at the global level but vanishes within subsets of genes with similar expression levels. The only remaining pattern is seen in the low expression quantile. Therefore, the global pattern must stem from the fact that the groups of genes expressed at different developmental stages have different properties, in particular in terms of expression levels.

The proportion of highly expressed genes indeed varied drastically between developmental stages (Fig 5A-D). At the early stage (bud), the vast majority of genes belonged to the class with high or medium-high expression levels (70%), whose sequence evolves more slowly on average. In the intermediate stages, the proportion of highly expressed genes dropped (about 40%) and increased again at the last stage (about 60%). The proportion of low-expression genes followed of course a complementary pattern. Thus, the inverse hourglass pattern of conservation is partly driven by the fact that genes expressed at different stages of development have different levels of expression.

**Fig 5.**
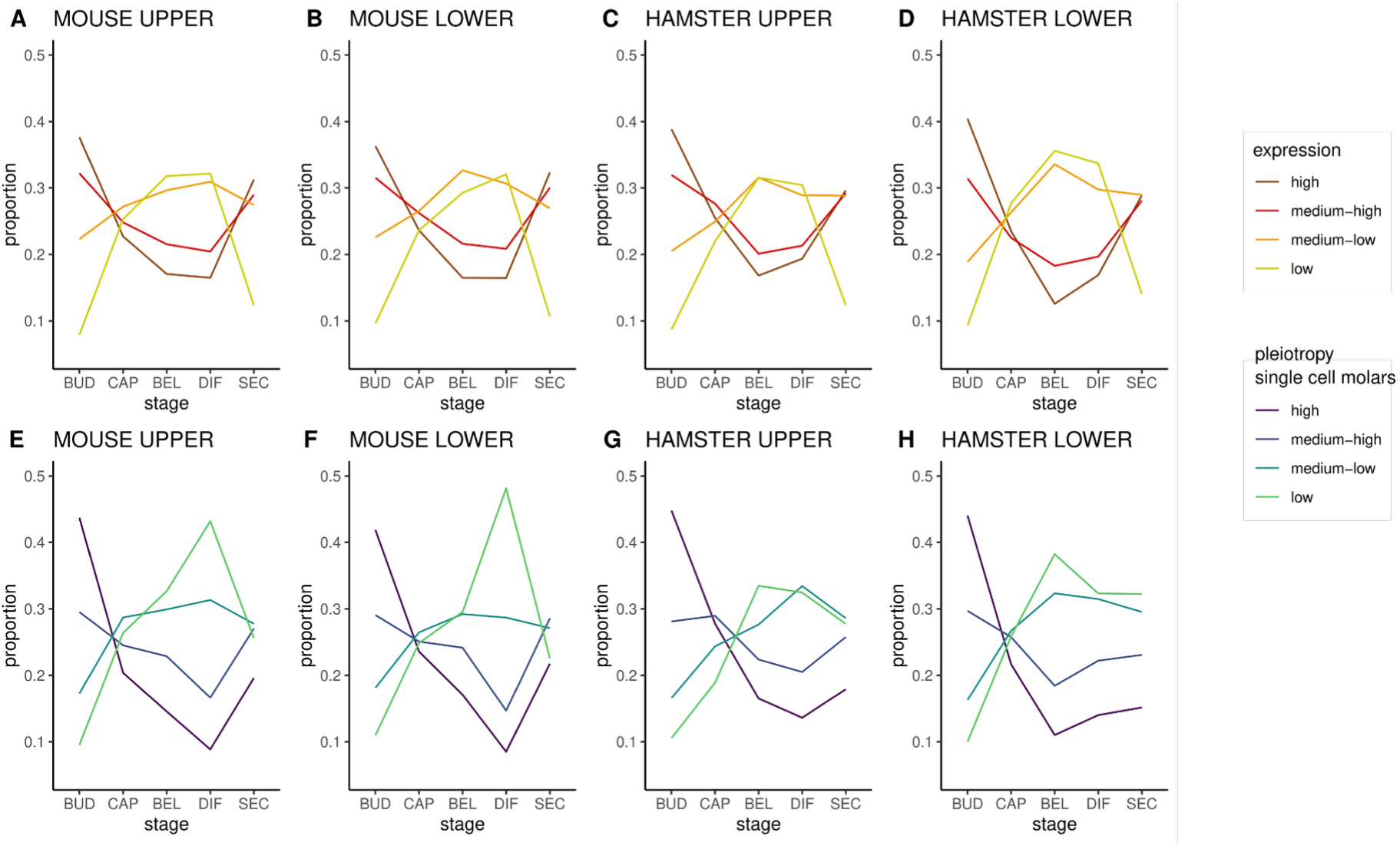
Genes expressed at different stages of development harbor different levels of expression and of cell expression specificity in molar. *A-D: Genes were split in quantiles according to their average expression level. The proportion of genes associated with each quantile is drawn for each developmental stage and for each molar. Quantile boundaries are computed separately for each molar, respectively quantile 1 to 4 in TPM for mus mx: 1.7, 20.9, 74.9,; for mus md: 1.6, 20.9, 74.8 for ham mx: 1.6, 17.2, 64.2; for ham md:1.5, 17.1, 63.9. E-H: Genes were split in quantiles according to their pleiotropy. The proportion of genes associated with each quantile is drawn for each developmental stage and for each molar. Quantile boundaries were computed on tau values measured on mouse molar single cell transcriptome, with three threshold values ranging from low to high pleiotropy (25% 0.37, 50% 0.61, 75% 0.85)*.

A same level of expression in a whole organ may be due to very high expression in a limited subset of cells, or to more moderate expression, but in a very large number of cells [35]. To assess this, we measured for each gene an index of expression specificity, the Tau index, based on the different cell types present in a developing tooth germ. We used published single-cell RNA-seq (sc-RNAseq) data for embryonic mouse lower molars, which we clustered in 23 cell populations [36]. We split genes in quantiles according to their level of cell-specificity and compared their proportions across tooth development (Fig 5E-H). At the bud stage, we observed that more than 40% of genes are pleiotropic (here: expressed in many cell types) and merely 10% are cell-specific. After that, the relative proportions reverse with about 10% pleiotropic genes at bell and differentiation stages. This mirrors the pattern observed for the quantiles of expression. In the latest stage however, the proportion of highly expressed genes increases again, while the proportion of highly pleiotropic genes remains modest. Hence, genes expressed in early and late stages tend to have different properties. Genes expressed in the early stage are expressed at a higher level in many cell types, while some genes expressed in the latest stage are also expressed at a high level but concentrated in fewer cell types (Fig S9).

Because expression specificity was estimated from a single scRNA-seq sample, it may not represent all cell types present in all stages of the molar timeserie. Incisors grow continuously in mice, therefore presenting in a single organ many of the tooth cell types and for these, spanning stem cell differentiation gradients. We measured cell expression specificity on another scRNA-seq dataset from adult mouse incisors and obtained similar results (Fig 6, 17 major cell subpopulations, [37]).

**Fig 6.**
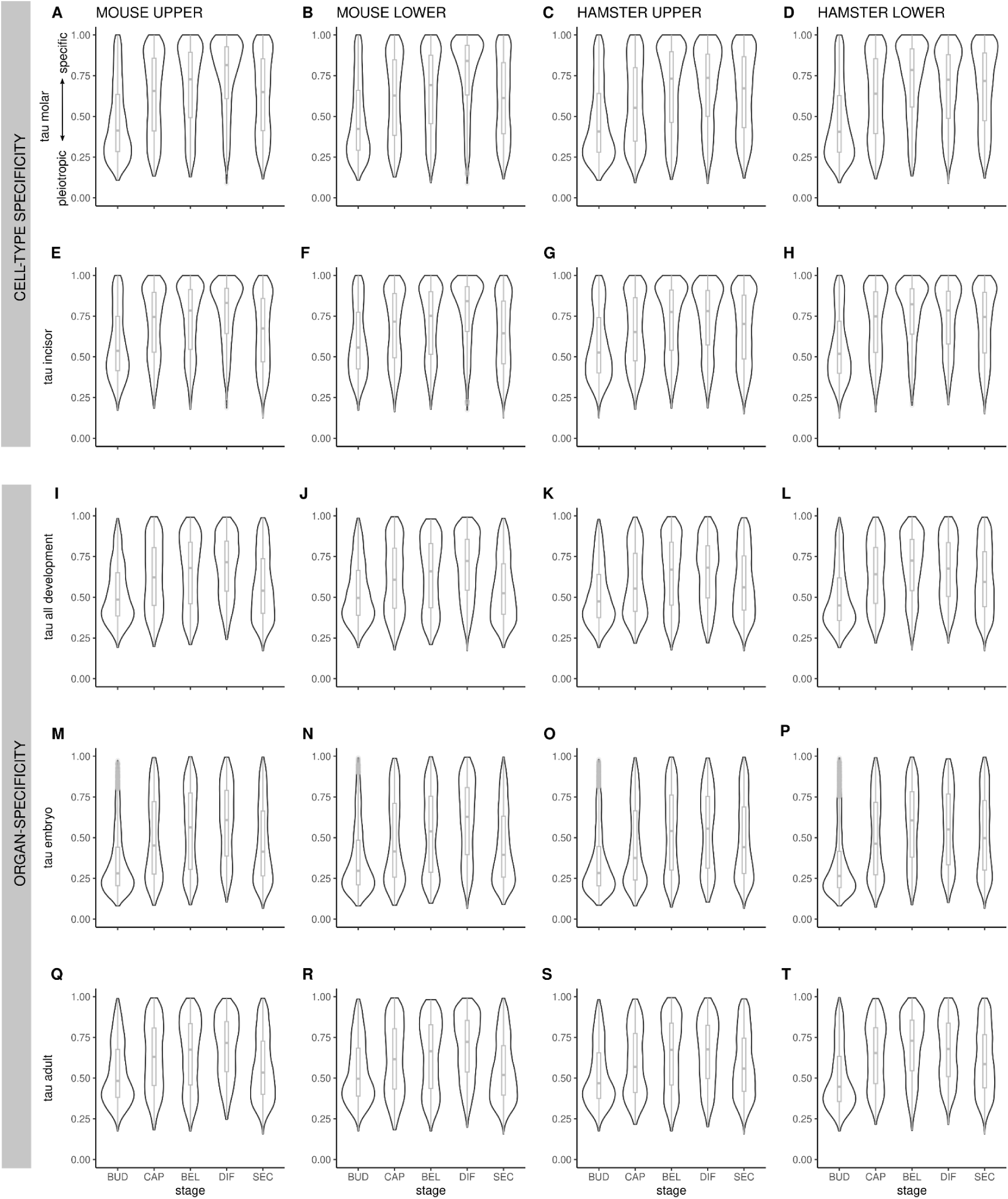
Distribution of expression specificity calculated by the tau index, in different data sets, for the 4 molars. *Cell expression specificity was measured on single cell RNA-seq data in mouse molar (A-D) and mouse incisor (E-H). Organ/stage expression specificity was measured on the Bgee data for all development (I-L), embryonic stages (M-P), and adult stages (Q-T). It is represented by a violin plot for each stage cluster. Boxplots represent the median and the lower 25% and top 75% quantiles. Tau values range from 0 (pleiotropic) to 1 (organ/cell specific)*.

**Fig 7.**
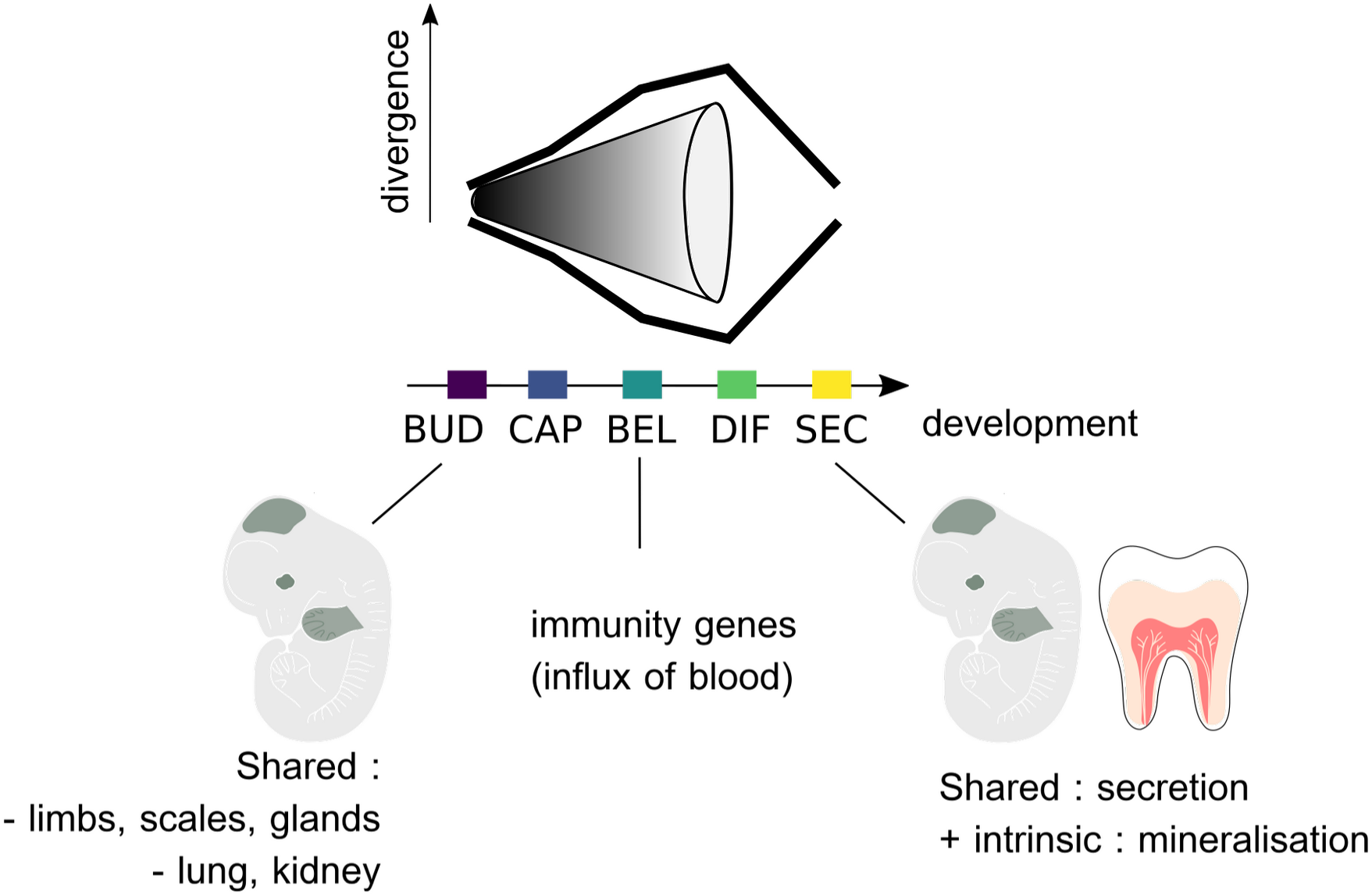
The conservation of the expression and sequence of genes expressed in molar development forms an inverted hourglass pattern, attributable to processes shared with other organs and intrinsic to the tooth.

Finally, we also computed tissue-specificity based on a meta-analysis of a large number of organs present in the Bgee database [38]. We found similar results to those from the molar data, with genes in coexpression clusters peaking at early and late stages being more pleiotropic (here: expressed in many stages and organs) than genes peaking at mid stages (Fig 6).

## DISCUSSION

While hourglass patterns of gene evolution have been found in many species and organs, rodent molar development appears characterized by an inverse hourglass which has been rarely seen (but see [16,39–42].

One of the main patterns in the field of evolution and development, which has been found in many biological systems, is that the highest developmental constraints apply during mid-embryogenesis, creating a so-called hourglass pattern [43]. This is true for transcriptome similarities at the level of the whole embryo, in many phyla [40]. This is also true at the level of organs, since an hourglass pattern was also observed when comparing limbs and fins, with a conservation peak around E10.5 [18]. Another study encompassing several developing organs found that conservation decreases with time, which is also in accordance with the hourglass pattern, since their sampling starts later than E10.5 [10]. The general interpretation is that gene regulatory complexity is higher in mid-development, while early and late development are more permissive to the fixation of mutations.

In molars, we observed that expression conservation forms an inverse hourglass pattern, since transcriptome similarity decreases with time, before increasing again at the secretion stage. The fact that early molar development is more conserved than later stages is in accordance with previous studies sampling organogenesis, like in [10]. It also makes sense with respect to the nature and age of the processes involved. Many processes involved in early tooth bud morphogenesis are shared with other skin appendages such as limbs, scales or glands, but also with lung or kidney branching morphogenesis [44,45]. Teeth and scales are ancestral to all jawed vertebrates and limbs are ancestral to tetrapods, and the components of the underlying developmental programs represent a very ancient molecular toolkit [46,47]. This program is even further recycled in other instances in body development, as demonstrated for mammary gland in mammals or patagium outgrowth in bats and sugar gliders [48]. Such ancient homology and pleiotropy is an important source of evolutionary constraint, exposing these morphogenesis genes to stronger negative selection both on their expression in teeth and on their coding sequences.

But if molars shared the pattern of other organs, we would expect a “funnel” of decreasing evolutionary conservation over all their development. Instead, an inverse hourglass pattern is observed, which has rarely been described, with two finding between plant and animal embryos of different phyla [40], one being later criticized [49], one study of molting in drosophila [42], and two studies of plant inflorescences [16,39]. Interestingly, in both plant inflorescences and molars, mid-development involves the patterning of iterated structures in a growing bud, namely flowers in a growing meristem and cusps in a growing tooth bud. In both cases, heterochronies in this process were linked to differences in final morphology [16,24,39,50]. This opens the possibility that minimal transcriptome conservation is observed at mid-development in such systems, because more heterochronies are present there, in relation to morphological adaptation, and that these blur the conservation signal in a process that would otherwise be largely homologous. In agreement with this hypothesis, we observed that genes peaking at mid-development tend to transition to the neighboring stage more frequently.

However, while the transcriptomes of maize and sorghum inflorescences differ most at mid development, their coding sequences do harbor classic hourglass patterns. In molars, the inverse hourglass pattern is also observed in coding sequences.

The reasons for the final increase in conservation are unknown, but since it is not observed in other mammalian organs we hypothesize that this may be related to the very particular nature of dental development. We noticed that in the secretory stage, we find a more bimodal distribution of the pleiotropy index as compared to the bud stage, suggesting that two classes of genes are present: specific genes and highly pleiotropic genes. This stage is marked by ameloblasts and odontoblacts actively producing and secreting the extra-cellular matrix (ECM) and the mineral components which will form enamel and dentin (e.g. golgi and endoplasmic reticulum apparatus, vesicle and ion transport…). Although some proteins of the ECM are highly specific to teeth, most proteins involved in this cellular activity are likely to be highly pleiotropic. We believe this heavy use of conserved cellular machinery may be responsible for this bimodal distribution, and contribute to the final decrease of dN/dS.

As expected, the amount of positive selection on coding sequences peaks at the differentiation stage in mouse and plateaus in the bell/differentiation/secretion stage in hamster. Individual genes showing traces of positive selection at these stages are not particularly associated with known processes of molar development, which does not argue that they are associated with adaptive changes in tooth shape in rodents. On the contrary, they are enriched in immune-related genes, in particular the interferon pathway. The discovery of immune-related genes is common in analyses of positive selection, because these genes are at the forefront of host-pathogen interactions [51]. We believe that we find them in the late stages, because following the late cap stage, blood supply is set up and immune cells are recruited locally [52]. Although those immune cells and the control of inflammation in the pulp are surely important for tooth function [53], these cell types are also important elsewhere for organism immunity. In conclusion, the pattern of positive selection is largely driven by local recruitment of cells with pleiotropic functions.

The relationship between the rate of sequence evolution and the level of gene expression has been discussed for a long time [31,33,34,54–56]. We found that the intensity of purifying selection on coding sequences is stronger for highly expressed genes, genes with conserved temporal expression profiles, and genes expressed in many cell types/organs. These characteristics are often correlated with each other, for biological and technical reasons. By separating these effects, we show that the coding sequence of genes exhibiting expression peaks at early and late stages are more subject to purifying selection for different reasons. Genes whose expression peaks in early molar development are expressed in more tooth cell types and in more adult and developmental organs. This higher pleiotropy is consistent with the fact they are used in developmental processes recycled many times in the organism (see above). Pleiotropy puts constraints on the evolution of coding sequences and has been linked before with periods of conservation of development [8,10]. By contrast, genes whose expression peaks in late molar development have high intra-cellular levels of expression, and are not as pleiotropic. Several models have been proposed to explain why highly expressed genes evolve slowly, including mistranslation, protein misfolding, and protein misinteractions [32,33,54]. This could apply to stages where genes necessary for enamel and dentin formation start being highly expressed.

In conclusion, in molars, different types of genes are used at the different stages of molar development in terms of intracellular expression level and in terms of pleiotropy. This reflects the colonization and differentiation of specific cell types and of the activation of specific developmental processes. Hence, the inverse hourglass pattern of conservation is both driven by the evolution of developmental processes intrinsic to teeth and by negative and positive selection essentially extrinsic to the teeth, via the level of gene pleiotropy. Similar factors likely underlie the patterns observed in other systems. In depth evolutionary analyses of organ development at the single cell level [6] may help quantify and distantangle the effect of these different factors.

## METHODS

### Rodent breeding and embryo sampling

CD1 (CD1) adult mice and RjHan:AURA adult hamsters were purchased from Charles River (Italy) and Janvier (France) respectively. Females were mated overnight and the noon after morning detection of a vaginal plug or sperm, respectively, was indicated as ED0.5. Other breeding pairs were kept in a light-dark reversed cycle (12:00 midnight), so that the next day at 16:00 was considered as ED1.0.

Pregnant mouse females were killed by cervical dislocation. Hamster females were deeply anesthetized with a ketamine-xylasine mix administered intraperitoneally before being killed with pentobarbital administered intracardially. All embryos were harvested and thereby anesthetized on cooled Hank’s or DMEM advanced medium, weighted on a precision balance after excess of liquid was removed with a whatmann paper, as described in <u>(Peterka et al. 2002)</u> and immediately decapitated.

### Ethics Statement

This study was performed in strict accordance with the European guidelines 2010/63/UE and was approved by the Animal Experimentation Ethics Committee CECCAPP #2018022617168148 (Lyon, France).

### Sample preparation

A total of 103 molar samples were obtained, corresponding to upper and lower molars from staged embryos in hamster and mouse. 36 samples were collected specifically for this study to complete our previously published data [24] Table S1. They cover in duplicates the whole period of tooth development in mouse from embryonic days (E) E12.5 to E22.5, equivalent to postnatal (PN) PN2 and in hamster (from E11 to E19.5, equivalent to PN2). Each sample contains two whole tooth germs, the left and right first molars (M1) of the same female individual, and for a given stage, upper and lower samples were prepared from the same individual. Dissections were as tight as possible to include only M1 tissue, excluding the posteriormost molar prospective regions (as in [50]). The heads of harvested embryos were kept for a minimal amount of time in cooled advanced DMEM medium (small scale) or advanced DMEM medium (large scale). The M1 lower and upper germs were dissected under a stereomicroscope and stored in 200-500µl of RNA later (SIGMA), adjusting for sample size. Total RNA was prepared using the RNeasy micro kit from QIAGEN following lysis with a Precellys homogenizer. RNA integrity was controlled on a Bioanalyzer or Tapestation (Agilent Technologies, a RIN of 10 was reached for all samples). PolyA+libraries were prepared with Illumina stranded mRNA prep kit), starting with 150 ng total RNA as in the previous study, and sequenced on an Illumina Hi-seq4000 sequencer at the Lausanne Genomic Technologies Facility. Because previous libraries had been generated with the non-stranded Illumina protocol and sequenced with a different design and platform, we also included 4 technical replicates, one per each end of the timeseries. We obtained on average 44 millions reads per sample with 150bp single-end strand-specific data for 40 samples (batch 2 and 2.1 for technical replicates, Table S1) and 100bp paired-end non-strand specific data for 65 samples (batch 1). Raw data are publically available in ENA with project accession numbers PRJEB52633 and PRJEB84925.

### Expression levels

These reads were mapped by using STAR (version 2.7.3a [57]) to reference sequences for golden hamster and house mouse transcriptomes. To generate them, we retrieved mouse and hamster genomes and annotations from Ensembl (release 98, January 2020, assemblies GRCm38 and MesAur1.0, [58]). The number of reads per genes was obtained by STAR –quantMode GeneCounts option, with default settings accounting for the strand-specificity of the data. The mapping rate is slightly higher in mouse than in hamster, with on average 80% of reads mapping unambiguously to annotated genes in mouse, and 65% in hamster. A table of counts and of TPM (transcripts per millions) values were created by a custom script based on these raw counts and on transcript length taken from Ensembl (release 98).

### Orthology relationships

Pairs of one to one orthologs between mouse and golden hamster were retrieved through Ensembl (release 98) by using biomaRt R library (version 2.48.0, [59]). Among these genes, only those with a MGI identifier were kept for further analysis. We kept 15910 pairs of orthologs.

### Multivariate analyses

The total table of raw counts contained 11342 genes with 1:1 orthologs in mouse and hamster, and with expression data in tooth development. We removed the samples corresponding to the upper and lower molars of one outlier individual (hamster E11.5), retaining 103 samples. This dataset was first normalized using DESeq2 with the sequencing replicate as a batch effect (3.11, [60]). We implemented PCA using the prcomp function from the stats package. Between-class analyses were used to estimate the effect of different factors on the PCA axes [61].

### Clustering of coexpressed genes

Stage-specific genes: Bgee Call version 1.4.0 was used with default parameters to generate present/absent gene expression calls [62]. Ensembl (release 84) was used for the mouse reference intergenic sequences, and community reference intergenic regions were used for the hamster.

Clusters of coexpressed genes: For each tooth type, the corresponding table of raw counts was first normalized with DESeq2 with the sequencing replicate as a batch effect (3.11, [60]). EISA clustering (eisa 1.44.0, [63]) was made with random starting clusters of samples (“random seeds”) and guided starting clusters of samples (“smart.seeds”). In the later case, starting conditions were obtained by grouping samples per developmental stage (bud, cap, bell, differentiation, secretion). The function ISAIterate was used to optimize the samples and genes grouping into clusters with the following parameters: thr.feat=1, thr.samp=1, convergence=“cor”. In the output, each gene has a score for each module. We assigned a gene to a particular module if the score of this gene is maximum in this module. Genes with a null score in all modules were not assigned.

### Functional enrichment

Functional enrichment was performed with the clusterProfiler [64] and ReactomePA [65] packages, using the enrichGO function to compute enrichment analyses in biological processes (BP) and molecular functions (MF) and the enrichPathway function to perform pathway enrichment analysis. All analyses were performed with p.adjust < 0.1 and a maximum of 50 categories were reported. terms were sorted by fold enrichment computed as following: FoldEnrichment = GeneRatio / BgRatio, with GeneRatio = k/n and BgRatio=M/N; with n genes in a given coexpression cluster, M genes in the functional gene set considered, k the overlap between the coexpression cluster and the functional gene set and N the number of unique genes in the dataset.

### Rates of sequence evolution

Selective pressure on coding sequences, estimated by dN/dS between mouse and golden hamster, were retrieved for 14382 pairs of one-to-one orthologs through Ensembl (release 98) by using biomaRt R library (version 2.48.0).

Estimates of positive selection were extracted from the Selectome database (V7, based on Ensembl release 98, [66,67]). For each gene, and for a selection of branches of interest, genes with strong signal of positive selection (q-value < 0.2) were extracted and their functional enrichment was studied with StringDB [68]. We also obtained the likelihood ratios of models of H1 to Ho models with and without positive selection, from the branch-site model [69]. The obtained ratios, Δln*L,* represent the evidence for positive selection. We transformed Δln*L* with the fourth root to improve their normality, following [15,29,70]. We obtained data for 11,460 genes on the Muridae branch, 12,919 genes on the murinae branch, 13,257 genes on the *Mus* branch, and 10,936 genes on the Cricetinae branch.

We split genes according to their phase of expression, following the clusters defined by EISA. Then, we computed the mean of dN, the mean of dS, and took their ratio mean(dN)/mean(dS). We also directly took the mean and median dN/dS of the genes per cluster. To get an estimate of the uncertainty of these values, we bootraped gene contents by resampling genes with replacement within clusters 1000 times, and recomputed the mean/median dN/dS as above. The same procedure was used for the 4th root of likelihood ratios taken from selectome.

### Pleiotropy/tissue specificity

To estimate pleiotropy/tissue-specificity at a multi-organ and multi-stage scale, we extracted expression values for three datasets: all developmental stages, embryonic and post-embryonic mouse libraries from the Bgee database (version 15.2) in September 2024 [71]. We log transformed the expression values (TPM) and calculated the mean of the log transformed expression values per anatomical feature. We obtained data for 117 organs and 55,486 genes (all developmental stages); 38 organs and 55,410 genes (embryonic stage), 86 organs and 55,486 genes (post-embryonic).

To estimate pleiotropy in teeth, we extracted expression values from two published single-cell RNA-seq datasets from wild type and healthy mice, one in incisors and one in molars. In incisors, we obtained expression levels for 2889 cells and their annotations in 17 cell types (GSE146123, [37]. In molar, we obtained 30930 cells from E14.5 stage lower molars, annotated in 23 cell types (GSE142200, [36]). We used Seurat package to obtain pseudo bulk expression profiling for each cell type [72].

We estimated pleiotropy in these 5 datasets by calculating Tau as previously described [73]. This index takes into account the mean of the log transformed expression values and the number of organs considered. Tau was calculated as: τ = sum(1-xb)/(n-1) where n is the number of organs considered, and xb=x/xmax is the level of expression normalized by the maximum level of expression in the vector. τ ranges from 0 (ubiquitously expressed) to 1 (expressed in a single organ/developmental stage).

### Data and code availability

Raw data are publically available in ENA with project accession numbers: PRJEB52633 and PRJEB84925.

All custom code (run in R) used in this study is made available here: https://gitbio.ens-lyon.fr/LBMC/cigogne/molar_inverse_hourglass

## ACKNOWLEDGEMENTS

We acknowledge the contribution of several platforms of SFR Biosciences Gerland-Lyon Sud (UMS344/US8): the Plateau de Biologie Expérimentale de la Souris (PBES) (many thanks especially to Jean-Louis Thoumas, Tiphaine Dorel, Céline Angleraux, Marie Teixeira, Myriam Prudent), as well as the computer resources from CBPSMN (ENS Lyon).

We acknowledge the technical help of Anne Lambert, Alain Rubod, Mathilde Estevez-Villar, and the contribution of many students including Coraline Petit, Alice Lorenc, Margaux Pillon, Ludivine Rotard and Asma Benahmed.

## AUTHOR CONTRIBUTIONS

S.P., M.S. and M.R.R. conceived the study, acquired funding, designed and performed some experiments, supervised research and wrote the manuscript. M.M., M.E.V. and S.P. collected the embryos. J.G. performed the experiments, and prepared visualization. S.M and M.N. performed the experiments on Selectome and Bgee databases, respectively. All authors have read the manuscript.

